# Tandem duplication events in the expansion of the small heat shock protein gene family in *Solanum lycopersicum* (cv. Heinz 1706)

**DOI:** 10.1101/057182

**Authors:** Flavia J. Krsticevic, Débora P. Arce, Joaquín Ezpeleta, Elizabeth Tapia

**Affiliations:** CIFASIS, Centro InternacionalFranco Argentino de Ciencias de la Información y de Sistemas - CONICET - UNR - AMU, Rosario, Argentina; GADIB, Grupode Análisis, Desarrollos e Investigaciones Biomédicas, Facultad Regional San Nicolás, Universidad Tecnológica Nacional, San Nicols, Argentina; Cátedrade Genética, Facultad de Ciencias Agrarias, Universidad Nacional de Rosario, Zavalla, Argentina; Facultad de Ciencias Exactas, Ingeniería y Agrimensura, Universidad Nacional de Rosario, Rosario, Argentina

**Keywords:** sHSP, ripening, tomato, transcriptome, RNA-seq, tandem duplication

## Abstract

In plants, fruit maturation and oxidative stress can induce small heat shock protein (sHSP) synthesis to maintain cellular homeostasis. Although the tomato reference genome was published in 2012, the actual number and functionality of sHSP genes remain unknown. Using a transcriptomic (RNA-seq) and evolutionary genomic approach, putative sHSP genes in the *Solanum lycopersicum* (cv. Heinz 1706) genome are investigated. A sHSP gene family of 33 members is established. Remarkably, roughly half of the members of this family can be explained by nine independent tandem duplication events that evolutionarily determined their functional fates. Within a mitochondrial class subfamily, only one duplicated member, Solyc08g078700, retained its ancestral chaperone function, while the others, Solyc08g078710 and Solyc08g078720,likely degenerated under neutrality and lack ancestral chaperone function. Functional conservation occurred within a cytosolic class I subfamily, whose four members, Solyc06g076570, Solyc06g076560, Solyc06g076540 and Solyc06g076520, support ~57% of the total sHSP RNAm in the red ripe fruit. Sub-functionalization occurred within a new subfamily, whose two members, Solyc04g082720 and Solyc04g082740, show heterogeneous differential expression profiles during fruit ripening. These findings, involving the birth/death of some genes or the preferential/plastic expression of some others during fruit ripening, highlight the importance of tandem duplication events in the expansion of the sHSP gene family in the tomato genome. Despite its evolutionary diversity, the sHSP gene family in the tomato genome seems to be endowed with a core set of four homeostasis genes: Solyc05g014280, Solyc03g082420, Solyc11g020330 and Solyc06g076560, which appear to provide abaseline protection during both fruit ripening and heat shock stress in different tomato tissues.

## INTRODUCTION

Tomatoes are native to South America and thirteen species are currently known, including the ketchup-worthy commercial variety *Solanum lycopersicum*. The Solanaceae species are characterized by a high degree of phenotypic variation, ecological adaptability (from rainforests to deserts) and similar genomes and gene repertoires. Because of its commercial importance, *S. lycopersicum* (cv. Heinz 1706) is a centerpiece of the Solanaceae family. The completegenome of this species, comprising 950 Mb and ~35000 protein-coding genes, was released in 2012by the Tomato Genome Consortium. The small size of its diploid genome makes *S. lycopersicum* (cv. Heinz 1706) a good reference for thestudy of the Solanaceaespecies and explains the emerging use of this fruit as a model system for the study offleshy fruit development (Aoki *et al.* 2013). In this regard, it is worth mentioning that the Solanum lineage has experienced two consecutive genome triplication events and that these events have led to the noe- or sub-functionalization of genes controlling important fruit characteristics such as color and fleshiness (The Tomato Genome Consortium 2012). Historically,low molecular weight (12-40 kDa) chaperone-like proteins or small heat shock proteins(sHSPs) have been associated with stress tolerance factors by preventing the irreversible aggregation of misfolded proteins (Basha *et al.* 2015; Poulain *et al.* 2010). However, heat shock stress is not the only stimulus triggering sHSP gene expression and protein synthesis. Indeed, sHSP synthesis is also induced during fruit maturation (Low *et al.*2000;) Lawrence *et al.* 1997; Neta-Sharir *et al.* 2005) and certain development stages (Prasinos *et al.* 2005; Faurobert *et al*.2007) in both Arabidopsis and Solanaceae plants, suggesting the existence of acomplexchaperone-dependent regulating network associated with these processes to maintain cellular homeostasis. In fact, pre-genomic data on tomato sHSPs provides experimental evidence for at least 14 sHSPs (Frank *et al.* 2009;Lee *et al.* 2012; Baniwal *et al.*2004; (Sanmiya *et al.*2004). This number has almost doubled in the post-genomic era with the finding of around 26 sHSP genes responsive to several stress situations on different tissues, including heat shock stress on leaves (Fragkostefanakis *et al.* 2015) and microspores (Frank *et al.* 2009). We note, however, that similarly to other gene families in tomato (Andolfo *et al.* 2014, current annotation of the sHSP gene family may not be fully defined.Multiple-copy sHSP genes may have gone unnoticed due to intrinsic limitations of genome assembly software (Krsticevic *et al.*2010), and multiple sHSP genes may have been mis-annotated into a family of functionallyunrelated proteins known as ACD-like or HSP20-like (Bondino *et al.* 2012) based solely on the presence of a conserved alpha-crystallin domain IPR008978 (ACD or HSP20 domain). To uncover the actual size and organization ofthe sHSP gene family in the *S. lycopersicum* (cv. Heinz 1706)genome, transcriptomic data of putative sHSP genes is analyzed from an evolutionary perspective.

## MATERIALS AND METHODS

### Putative sHSP genes and transcriptome data in *S. lycopersicum* (cv. Heinz 1706)

A BlastP search against the Tomato protein database (ITAG2.4 Release, Sol Genomics Network) was first performed using the amino acid sequence of Solyc09g015020, characterized by a conserved ACD domain, as query. Aiming to capture all putative members of the sHSP gene family, every annotated protein containing the IPR008978 HSP20-like chaperone Interpro domain was also retrieved from the Sol Genomics Network database (ITAG release 2.40). In addition, every putative sequence related to sHSPs was retrieved from the Helmholtz-Muenchen tomato database using “small HSP” as search keyword. As a result, 58 putative sHSP sequences of *S. lycopersicum* (cv.Heinz 1706) were retrieved. Taking into account that nine *β*-sheets are expected in conserved ACD domains (Poulain *et al.* 2010), putative sHSP sequences were further characterized with the corresponding number of *β*-sheets. The ACD domain was identified with PROTEUS2, a web server supporting comprehensive protein structure prediction and structure-based annotation, http://wishart.biology.ualberta.ca/proteus2. (Montgomerie *et al.* 2008). Additionally, the number of *β*-sheets was estimated with Phyre2, a web server supporting the prediction of secondary structures, http://www.sbg.bio.ic.ac.uk/phyre2(Kelley *et al.* 2015).

Fruit ripening in tomato starts at the mature green (MG) stage, where an extensive metabolic reorganization takes place, evolves to the mature breaking (MB) stage and, ten days later, reaches the mature red (MR) stage. Illumina RNA-seq read files of *S. lycopersicum* (cv. Heinz 1706) from two biological replicates of the MG,MB and MR fruit ripening stages were downloaded from the DDBJ Sequence Read Archive (DRA) database (http://trace.ddbj.nig.ac.jp/dra/index_e.html.) Concerning complementary analysis of putative sHSP genes during the growth stages of tomato fruit, RNA-seq read files from 1 cm and 2 cm immature green fruit were retrieved. Finally, concerning complementary tissue preferential analysis of putative sHSP genes, Illumina RNA-seq read files from leaf, root, flower and flower bud were also retrieved (see Table S1). SRA files were converted to FASTQ files using the fastq-dump utility of the SRA toolkit. *S. lycopersicum* (cv. Heinz 1706). RNA-seq data from the set of MG, MB and MR fruit maturation stages, the 1cm and 2cm immature stages, and the root, stem and leaf plant tissues were then aligned back to the tomato genome assembly SL2.50 with annotation ITAG2.40 (The Tomato Genome Consortium 2012) using TOPHAT version 2.0.13 (Trapnell *et al.* 2009) with bowtie2 version 2.2.4 and default settings. Read counts for each gene were quantified using coverageBed from bedtools version 2.25.0. Fruit maturation entails a development process that may cause varying mRNA levels between group samples, which may discourage the use of standard normalization methods (Aanes *et al.* 2014). To shed some light on this issue, the quantroStat statistic was used (Hicks and Irizarry 2015). Differences between ripening groups were detected at the *α* = 0.01 significance level. In this scenario, one must still decide if detected differences are likely to be biologically or technically driven, in which case standard normalization methods should be applied. Taking into account that only two biological replicates per group were available, which may render the quantroStat imprecise, and that relative logarithmic expressions (RLE) box plots across groups showed an almost in line appearance, the Trimmed Mean of M-values (TMM) normalization method in the edgeR Bioconductor package (Robinson *et al.* 2010) version 3.12.0 was applied. In any case, normalization factors close to unity were obtained, suggesting that the use TMM normalization to remove technical variability would have minimal impact on downstream data analysis.

FPKM (Fragments Per Kilobase of transcript per Million mapped fragments) values were calculated to determine transcript abundance of individual genes. Average FPKM values were calculated for each pair of biological replicates for each sample (see Table S2).Following the criteria of previous studies (González-Porta *et al.*2013; Hebenstreit *et al.*2011), genes were considered expressed when average FPKM values were greater than or equal to 2. Differentialexpression between pairs of ripening stages was assessed with the exactTest function. The MG fruit stage was taken as baseline and edgeR’s exact test was applied to identify significant log_2_ fold changes (FC) of the MB and MR stages relative to the MG baseline at the *α* = 0.01 significance level (see Figure 1). Positive log_2_ FC values indicated up-regulation, negative ones indicated down-regulation and zero indicated constant gene expression relative to the MG baseline. In these studies, the *tip41* housekeeping gene (Solyc01g107420) was used as negative control for differential gene expression while the *hsp70* gene (Solyc11g020040) was used as a positive control for up-regulation during fruit ripening (Exposito-Rodriguez *et al.* 2008); (Fei *et al.*2004).

### Assessing the phylogeny of putative sHSP genes in *S. lycopersicum* (cv. Heinz 1706) and evolutionary times of duplication events

A phylogeny-based functional analysis was used to complete (Eisen *et al.* 1998) preliminary chaperone function prediction obtained through RNA-seq expression analysis of putative sHSPs in *S. lycopersicum* (cv. Heinz 1706). 11 sHSP sequences of *Arabidopsis thaliana* (Waters *et al.* 2008), known to be representative and conserved between angiosperms, together with the 58 putative sHSP sequences of tomato,were used to construct a initial phylogenetictree. Additionally, the subcellular localization of putative sHSPs was obtained from analogous clusters formed in the phylogenetic tree (see Figure S1). Molecular evolutionary analyses were conducted with MEGA version 6 (Tamura *et al.* 2013). Amino acid sequences were aligned using ClustalW with default settings and a Maximum Likelihood analysis based on the Jones-Taylor-Thornton (JTT) substitution model for proteins with a gamma distribution was performed (Larkin *et al.* 2007). Acondensed tree was built with a bootstrap method set to work with 100 replications anda cut-off value of70% (Hillis and Bull 1993). Aiming to evidence evolutionary relationships in the resulting sHSP gene family, a second phylogenetic tree was built with curated sHSPs obtained after joint RNA-seq and phylogeny-based functional analysis. In this case, a Maximum Likelihood analysis based on a WAG substitution model for proteins with a gamma distribution was used.

**Figure 1.**
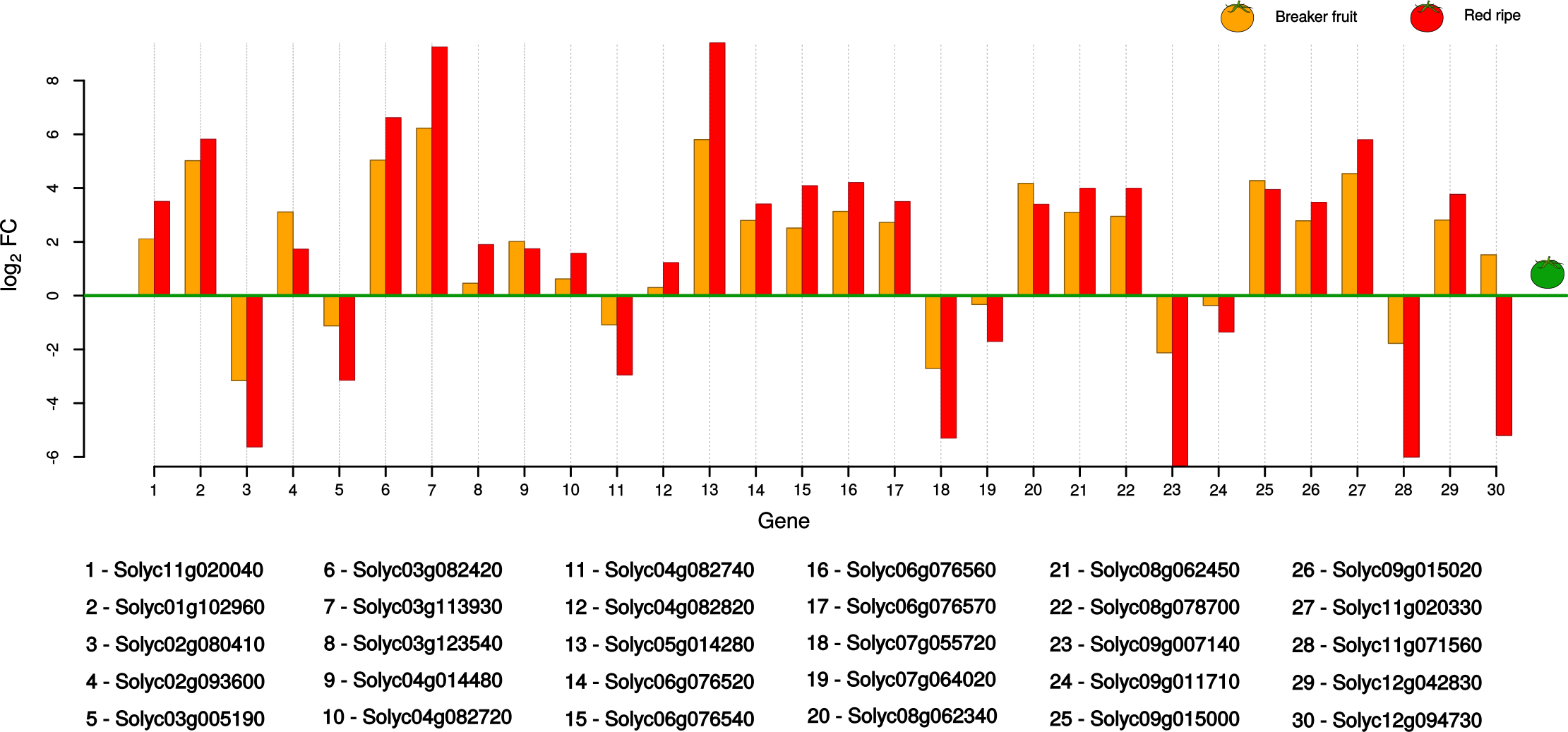
Putative sHSP genes differentially expressed during tomato fruit ripening (29 out of a total of 58). Differential expression at the mature breaker (MB) and mature red (MR) fruit ripening stages relative to the reference mature green (MG) one is quantified as log2 FC. The hsp70 gene or Solyc11g020040 (#1) is used as a positive control of up-regulation.

Following Xia *et al.* (2011), the DnaSP tool (Librado and Rozas 2009) was used for the computation of occurrence time *T* of the tandem duplicated genes. Briefly, DnaSP provides the mean number *K*_*s*_ of synonymous substitutions per synonymous site between pairs of duplicated genes. The occurrence time of duplication events can be calculated using 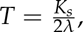 where λ is the clockwise substitution rate, a characteristic parameter of each species. Based on the work of Yang *et al.* (2008), λ = 1.5 10^-8^ of *A. thaliana* was used to find the approximate value of *T* in tomat.

## RESULTS

### Putative small heat shock proteins (sHSPs) in the *S. lycopersicum*(cv. Heinz 1706) genome

To begin the reconstruction of a large-scale picture of the sHSP family organization in the *S. lycopersicum* (cv. Heinz 1706) genome, 58 putative sHSPs were mapped to their chromosomes. Putative sHSP genes were found to be evenly distributed (Pearson’s Chi-squared test based on 10^7^ replicates, p-value=0.5023) across the 12 tomato chromosomes. Despite this random distribution, an integral analysis of the expression profiles of putative sHSPs genes across multiple conditions and tissues together with their phylogenetic relationships may help define and characterize the sHSP gene family.

### Differential expression of putative sHSP genes during tomato fruit ripening

Differential expression analysis pointed out 29 out of the initial set of 58 putative sHSP genes as differentially expressed in the MR fruit ripening stage relative to the reference MG one see (Figure 1). A sub groupof 20 putative sHSP genes was found to be up-regulated during the fruit ripening process,with expression intensity rising from MG to MR, suggesting that these sequences were indeed sHSPs. On the other hand, a subgroup of nine putative sHSP genes was found to be down-regulated in the MR stage relative to the reference MG one. This pattern of down-regulation may be caused by two non-mutually exclusive events: natural gene expression declination during fruit senescence or the fact that putative down-regulated sHSP genes are actually HSP20-like genes. To further support the first evidence of chaperone functionality in putative sHSP genes being up-regulated in the MR stage relative to reference MG one, and to clarify the status of those appearing as down-regulated, not differentially expressed or even not expressed, a phylogenomic analysis was performed.

### Phylogeny-based annotation of putative sHSP genes

A set of 20 clusters representing the four major monophy-logenetic clades characteristic of higher plants, MCI, MCII, MCIII and MCIV (Bondino *et al.* 2012), was identified in a phylogenetic tree built from 58 putative sHSPs sequences in *S. lycopersicum* (cv. Heinz 1706) and 11 sHSP reference sequences in *A. thaliana* (see Figure S1s).

A set of 18 sHSP genes distributed in 10 clusters, containing 16 up-and two down-regulated members during fruit ripening, was clearly identified based on functional orthology with *A. thaliana* and previous experimental evidence. Another set of 13 sHSP genes distributed in four clusters (#14, #13, #9 and #7) was identified after careful analysis of previous experimental evidence, completeness of the ACD domain and presence of other domains (Table S3). Cluster #14 initially had two paralogous members, Solyc08g078710 and Solyc08g078720, map together with singleton Solyc08g078700. The three sequences show around 47% sequence identity, suggesting a common origin product of a tandem duplication event (see Figure S3). Altogether, these evidences suggest Solyc08g078700 membership to Cluster #14. In addition, upregulated Solyc08g078700 is similar (46.7% identity and 63.8% similarity at the amino acid sequence level) to the mitochondrial 23.6 kDa Q96331 protein in *A. thaliana,* already classified as a sHSP (Scharf *et al.* 2001). Altogether, these evidences suggest that, although Solyc08g078710 is not expressed during fruitripening and Solyc08g078720 is not differentially expressed, both are sHSPs. Cluster #13 contains two paralogous members, Solyc10g086680 and Solyc09g011710. Although Solyc10g086680 is not expressed during fruit ripening and Solyc09g011710 is down-regulated, they both have a complete ACD domain and, additionally, Solyc09g011710 has been reported to be down-regulated under heat shock stress in leaves (Fragkostefanakis *et al.* 2015). Altogether, these evidences indicate that both Solyc10g086680 and Solyc09g011710 are sHSPs. Cluster #9 contains four paralogous members, Solyc01g009200 and Solyc01g009220, which are not expressed during fruit ripening but are down-regulated during fruit development (see Figure S2, and Solyc11g071560 and Solyc09g007140, which are down-regulated during fruit ripening. Taking into account that all members of this cluster but Solyc01g009220 have a complete ACD domain, the evidences indicate that the four members of Cluster #9 are likely sHSPs (see Table S3). Cluster #7 contains four paralogous members, Solyc04g082720 which is up-regulated during fruit ripening, Solyc04g082740 which is down-regulated, Solyc01g098810 which is not differentially expressed and Solyc01g098790 which is not expressed. In addition, all members of this cluster but Solyc01g098790 have a complete ACD domain. Altogether, these sources of evidence indicate that the four members of Cluster #7 are sHSPs. Note, however, that Cluster #7 has a strikingexpression pattern, with both up- and down-regulated members during tomato fruit ripening (see discussion below). In agreement with the joint presence of the ARID and ACD domains (Riechmann *et al.* 2000), a set of 6 putative sHSPs clustering together in Cluster #4 (Figure S1) turned out to be transcription factors, i.e., HSP20-like genes. While two of these transcription factors have been reported previously (Bondino *et al.* 2012), four of them could be new. Similarly, another set of 14 putative sHSPs distributed in 5 clusters are also potential HSP20-like genes since they all lack functional and evolutionary evidence to support sHSP family membership. Finally, two sHSP genes, Solyc02g093600 and Solyc04g072250, were identified in the set of seven putative sHSP genes that remained unclustered. Solyc02g093600 is up-regulated during fruit ripening, has a complete ACD domain and previous experimental evidence(Fragkostefanakis *et al.* 2015). On the other hand, structural features of Solyc04g072250, including the presence of an ORF, absence of a premature stop codon and a complete ACD domain, suggest its chaperone functionality. However, previous experimental evidence (Fragkostefanakis *et al.* 2015) and lack of expression during fruit ripening suggest that Solyc04g072250 is actually non-functional. Based on the these considerations, a sHSP gene family of around 33 members can be defined in the *S. lycopersycum* (cv. Heinz 1706) genome. Regarding the fruit ripening process, this result brings new experimental evidence about the chaperone function of four members of the sHSP gene family, Solyc04g082740, Solyc09g007140 and Solyc11g071560, which appear down-regulated, and Solyc04g082720, which appears up-regulated. Additionally, we extend the chaperone function of Solyc03g123540 and Solyc02g093600, already reported as sensitive to heat shock stress, to fruit ripening.

### Subcellular localization of sHSP genes

Subcellular localization of proteins can provide important evidence about their function (Emanuelsson *et al.*2000). Based on evolutionary relationships andin agreement with previous experimental evidence (Figure S1), 22 sHSPs are predicted to be ubiquitously distributed across cellular compartments and organelles Table S4. Briefly, 11 of them are predicted for the cytoplasm (CI), two for chloroplasts (CP), three for the endoplasmic reticulum (ER), five for the mitochondria (MT)and one for the peroxisoma (PX). Two sHSP genes, Solyc05g014280 and Solyc03g113930,whose products are presumed for the CP and ER, show the highest levels of differentialexpression during fruit ripening (Table S5). Remarkably, four sHSP genes found to be highly responsive to fruit ripening stress, Solyc05g014280, Solyc03g082420, Solyc11g020330 and Solyc06g076560, whose products are presumed for the CP, ER and CI, have been also reported to be highly responsive to heat shock stress in leaves (Fragkostefanakis *et al.* 2015) and microspores (Frank *et al.* 2009), suggesting the existence of a core set of sHSP genes important to maintain cellular homeostasis under 1, 4, 6, 8 and 9 (both stress situations (Figure 2).

**Figure 2.**
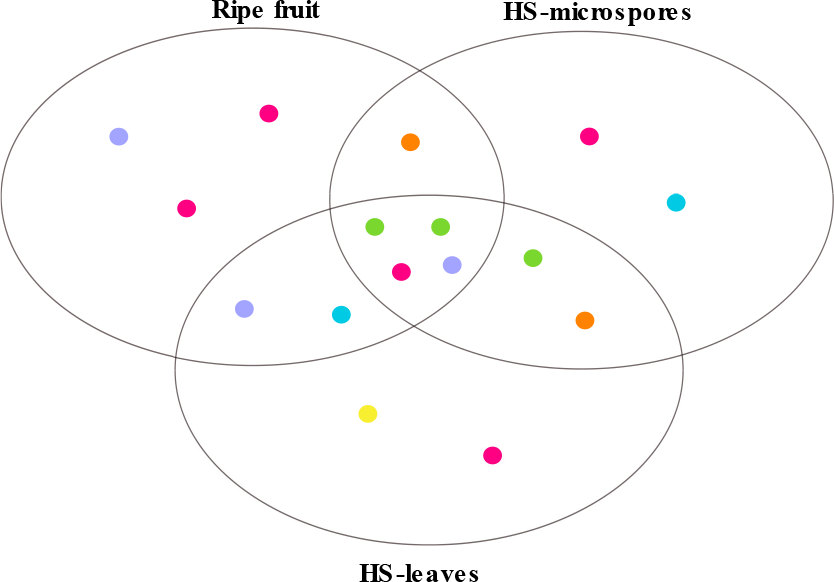
Top 10 sHSP genes responsive to fruit ripening and heat-shock stress in leaves and microspores. For each condition,sHSP genes targeted to the endoplasmatic reticulum (ER), the cytosolic classes I and II (CI, CIsIII and CII), perixoma (PX), the chloroplast (CP) and the mitochondrial (MT) are shown. Four sHSP genes, Solyc05g014280, Solyc03g082420, Solyc11g020330 and Solyc06g076560, targeted for the CP, the ER and theCI, are responsive in all conditions.

### Tandem duplication events in the sHSP gene family

Tandem and segmental duplication are main sources of diversity for the evolution of large gene families in plants (Cannon *et al.* 2004). A phylogenetic analysis of the sHSP gene family in *S. lycopersicum* (cv. Heinz 1706) revealed 17 sHSP genes (~51%) produced by tandem duplications events (see Figure 3). These genes are organized into six subfamilies that map to chromosomes 1,4, 6, 8 and 9. Certainly, segmental duplication has also contributed to the expansion of gene families in plants. However, its role may be less pronounced inthe diversification of the sHSP family (Waters *et al.* 2008). To shed light on this issue, duplications of sHSP genes were investigated with the MCSCAN too(Tang*et al.*2008) and little evidence about a dominant role of segmental duplication in *S. lycopersicum* was found. Duplication analysis basedon the identification of synteny blocks showed only 2 segmental duplications among chromosomes 6, 9 and10 involving three genes, Solyc06g076520, Solyc09g011710 and Solyc10g086680. These segmental duplications may be attributable to the last whole-genome triplication (91 - 52Myr) occurred in the Solanum lineage (The Tomato Genome Consortium 2012).

Three sHSP subfamilies are useful to describe the alternative functional outcomes of tandem duplicated sHSP genes in *S. lycopersicum* (cv. Heinz 1706). Afirst subfamily involves three MT class sHSP genes mapping together to a ~11.4 kb region in chromosome 8 (SL2.40ch08:59625875.59637274). Notably, in this subfamily,only the basal gene Solyc08g078700 appears as clearly functional, while the other two subfamily members, Solyc08g078710 and Solyc08g078720, seem to be losing their ancestral chaperone function. A second subfamily involves four functional intronless CI class sHSP genes mapping together to a ~17.9 kbp region in chromosome 6 (SL2.40ch6:47.547k.47.564k). Three members of this subfamily, Solyc06g076540, Solyc06g076560 and Solyc06g076570, have been previously reported by (Goyal *et al.* 2012) in *S. lycopersicum* (cv. Ohio 8245). Now, a fourtmember, Solyc06g076520, is reported. Notably, the four members of this subfamily support ~57% of the sHSP transcripts in the MR fruit ripening stage (Table S2). Furthermore, subfamily members Solyc06g076540 and Solyc06g076560 are among the most differentially expressed sHSP genes during fruit ripening (see Table S5). Finally, a third subfamily involves two sHSP cytosolic/nuclear genes, Solyc04g082720 and Solyc04g082740, mapping together to a ~9.1 kb region in chromosome 4 (SL2.50ch04:66300179.66309278). Notably, although both members of this subfamily are functional, their temporal expression patterns over development and ripening suggest that they are undergoing a sub-functionalization process.

Identification of multiple-copy genes in tomato, like the one presented here for the sHSP gene family, can contribute to reduce the uncertainty of estimations about exploitable phenotypic variation, which could be considerably useful in commercial tomato breeding programs.

## DISCUSSION

### Small heat shock proteins (sHSPs) in the *S. lycopersicum* (cv. Heinz 1706) genome

Even with the large amount of genomic data now available, the number and functionality of sHSP genes in the Solanaceae family remain largely unknown, and their functionalannotation is often inconsistent across authors and databases ( see Table S3. An evolutionary perspective on the transcriptome analysis of *S. lycopersicum*(cv. Heinz 1706) allowed us to define a sHSP gene family of around 33 members. Families of sHSP genes in plant species tend to be rather large and variable in size: 19 sHSP genes have been reported in *A. thaliana* (Scharf *et al.* 2001; Siddique *et al.* 2008), 39 in rice *Oryza sativa* (Ouyang *et al.* 2009) and 51 in *Glycine max* (Lopes-Caitar *et al.* 2013). Despite this variability, the proportion of sHSP genes in plant genomes is roughly constant, ranging approximately from 0.06 to 0.1%. The proportion of sHSP genes in *S. lycopersicum* (cv. Heinz 1706), 0.095% or 33 out of a total of 34727, is in accordance with these previous studies, suggesting that the totality of members of the sHSP gene family has been uncovered in tomato. Note, however, that the actual number and location of sHSP genes in the ~7000 domesticated lines of *S. lycopersicum* collected in the EU-SOL BreeDB database (https://www.eu-sol.wur.nl) may vary according to directional selection pressures (Ercolano *et al.* 2014).

### Tandem duplication events and the expansion of the sHSP gene family in tomato

The main function of sHSPs is to maintain the homeostasis of cellular proteins. The importance of this ubiquitous function supports the presence of redundant sHSPs so that if one of them fails, the others are ready to supply their chaperone function. Evolutionary forces clearly affected and modeled the sHSP gene family (Ohno 1970). Roughly half (17) of the sHSP genes in the *S. lycopersicum* (cv. Heinz 1706) genome can be explained by tandem duplication events. In most of these events, redundancy tends to be eliminated so that one of the copies retains its ancestral function while the other becomes a pseudogene (Zheng and Gerstein 2007).

Neutral evolutionary processes seem to be a valid argument to explain the behavior of two of three MT class tandem duplicated sHSP genes, Solyc08g078700, Solyc08g078710 and Solyc08g078720, mapping together to a 11.4 kb region in chromosome 8. While the basal Solyc08g078700 gene retained its ancestral chaperone function and evolved under purifying selection (see Figure S3 and associated Table), its two accompanying copies degenerated, Solyc08g078710 and Solyc08g078720, Functional redundancy also seems to a bea valid possibly under the effect of neutrality. Although Solyc08g078710 has a complete ACD domain, it is expressed neither in plant tissues (leaf, root, flower and flower bud) nor during fruit development (1 cm and 2 cm), fruit ripening or heat shock stress, probably due to variations in the promoter architecture of the 5’UTR region. Conversely, although Solyc08g078720 is expressed in all plant tissues, it is insensitive to fruit development, fruit ripening or heat shock stresses, probably due to the presence of an incomplete ACD domain with only seven beta sheets (see Table S3). Altogether, these evidences suggest that both Solyc08g078710 and Solyc08g078720 did not retain their ancestral chaperone function. Functional redundancy seems to be to a be a valid argument to explain the behavior of four Class I tandem duplicated intronless sHSP genes, Solyc06g076520, Solyc06g076540, Solyc06g076560 and Solyc06g076570, mapping togethertoa ~17.9 kb region in chromosome 6 (SL2.40ch6:47.547k.47.564k). If there is abiological reason for this sHSP gene subfamily to stay in array in a chromosome 6 region, e.g.due to its important relative contribution to differential expression and transcriptabundance of sHSP genes during fruit ripening, a high degree of conservation of this subfamily across close Solanum species should be expected. In effect, duplication analysis suggests that only Solyc06g076520 originated during the last whole-genome triplication in the Solanum lineage (together with Solyc09g011710 and Solyc10g086680 in Cluster#13). The remaining members of Cluster #2, Solyc06g076570, Solyc06g076540 and Solyc06g076560, seem to be the product of tandem duplication events, the first of which took place ~13 Myr ago (Figure S4). Taking this together with collinearity results between potato and tomato at the chromosome 6 region of Cluster #2, we can hypothesize that gene associations in Cluster #2 have indeed been maintained, thus reflecting their importance in the sHSP gene family. We note, however, that orthologous genes of Solyc06g076560 are absent in the *Solanum tuberosum* and *Solanumpennelli* genomes. Actually, Solyc06g076560 is a paralogous copy of parental Solyc06g076520 (97.9% nucleotide identity) suggesting the occurrence of a tandem duplication event exclusive to the *S. lycopersicum* clade (Figure 3).Finally, tandem duplicated genes may diversify and undergo some degree of neo-functionalization. In this regard, either of the copies may acquire a new beneficial function andthe other retain its ancestralfunction, or both copies may undergo sub-functionalization, with each copy being expressed uniquely at different tissues or with a temporal expression pattern (Lynch and Force 2000; He and Zhang 2005). In effect, temporal sub-functionalization seems to be a valid argument to explain the behavior of the two tandemduplicated sHSP genes, Solyc04g082720 and Solyc04g082740, mapping together toa ~9.1 kb region in chromosome 4. These sHSP genes show a complementary temporalexpression pattern (MacCarthy and Bergman 2007) during fruit development and ripening. According to the NexGenEx-Tom database(http://140.164.45.142/NexGenEx-Tom/expression/exp-search.aspx)Tom/expression/exp-search.aspx), while the peak of expression of Solyc04g082740 occurs during fruit development at the 3 cm of fruit size, the peak of expression of Solyc04g082720 occurs during ripening at the MR stage. Similarly to rice, where roughly half (19) of sHSP genes in the genome have been reported to be produced from tandem duplication events (Ouyang *et al.* 2009), and differently from *A. thaliana,* where only one tandem duplicated sHSP gene has been reported (Scharf *et al.* 2001), our results suggest that tandem duplication events have contributed largely to the expansion of sHSP gene family in *S. lycopersicum*.

**Figure 3.**
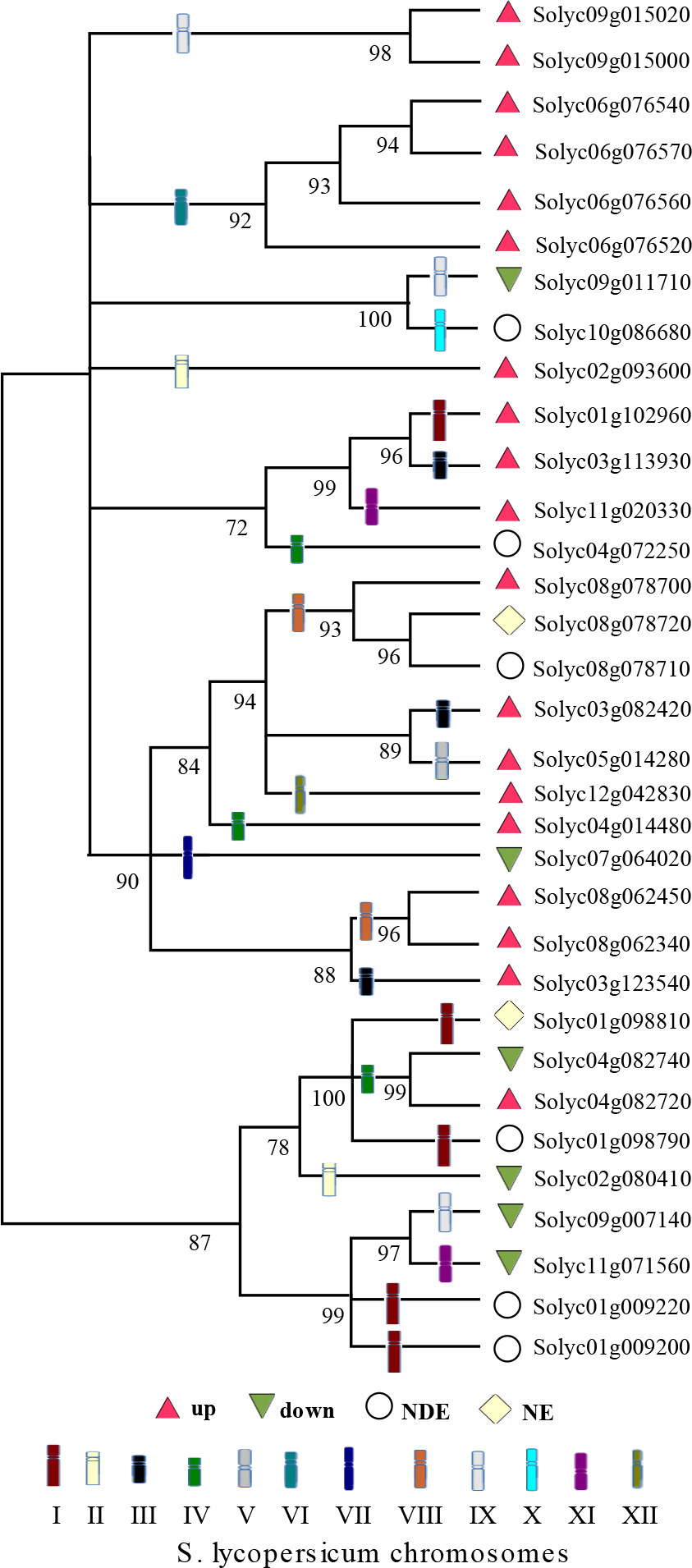
Phylogenetic relationships in the sHSP family of the *S. lycopersicum* (cv. Heinz 1706) genome. Amino acid sequences deduced from the 33 members of sHSP gene family are used. Gene expression profiles during fruit ripening are shown. Differential expression is measured at the MR stage relative to the MG reference one. Different symbols are used for depicting up-and downregulated, not differentially expressed (NDE) and not expressed (NE) genes. Aiming to highlight tandem duplication events thathappened during the evolutionary history of this family, the chromosome localization of sHSP genes is used for branch labeling. Nine tandem duplication events are present:one in chromosome 1 involving the pair Solyc01g009200 - Solyc01g009220, one in chromosome 4 involving the pair Solyc04g082720 - Solyc04g082740, two in chromosome 8 involvingthe pair Solyc08g062340 - Solyc08g062350 and the trio Solyc08g078700 - Solyc08g078710 - Solyc08g078720, three in chromosome 6 involving the quartet Solyc06g076520 - Solyc06g076540 - Solyc06g076560 - Solyc06g076570, and one in chromosome 9 involving the pair Solyc09g015000 - Solyc09g015020.

### A core set of homeostasis sHSP genes in tomato

Simultaneous analysis of most differentially expressed sHSP genes during fruit ripening and heat shock stress in leaves and microspores (Fragkostefanakis *et al.* 2015); (Frank *et al.* 2009) revealed the existence of four sHSP genes with a common and very sensitive response to both stress situations. Being sessile organisms,plants have evolved mechanisms to deal with and tolerate multiple stress situations. In contrast to other eukaryotic organism domains, plants are unique in expressing in a cell a multiplicity of cytosolic sHSPs and specific sHSPs targeted to the plastids, the endoplasmic reticulum and the mitochondria (Waters *et al.* 1996; Waters 1995; de Jong *et al.* 1998; Siddique *et al.* 2008).

Chloroplasts are responsible for photosynthesis as well as numerous other functionsin a plant cell (Jarvis and López-Juez 2013). During fruit ripening, a massive transformation process of chloroplasts into chromoplasts takes place and thus, repair and stabilization of proteins are required (Lawrence *et al.* 1997; Zeng *et al.*2014). A subfamilyof two chloroplast sHSP genes, Solyc03g082420 and Solyc05g014280, is present in the tomato genome. The Solyc05g014280 or *vis1* gene is highly differentially expressed during fruit ripening and seems to play a specific role in pectin depolymerization, a common event in fruit ripening, by reducing the thermaldenaturation of depoly-merizing enzymes in response to daytime elevated temperatures(Ramakrishna *et al.* 2003). On the other hand, the extensively characterized Solyc03g082420 or *hsp21* gene (Lawrence *et al.* 1997; Srivastava *et al.* 2010; Matas *et al.* 2011) is highly differentially expressed not only during fruit ripening and heat shock stress in leaves and plant microspores but also during plant development (Neta-Sharir *et al.*2005; Lambert *et al.* 2011). The endoplasmic reticulum (ER) plays a key role in a cell endomembrane system. It is involved in folding and assembling the majority of proteins that a cell secretes, in lipid synthesis, and insustaining cell homeostatic balance (Walter and Ron 2011; Howell 2013). In higher plants, endoplasmic reticulum sHSPs are accumulated in this compartment (Zhao *et al.* 2007) and may protect endoplasmic reticulum proteins from stress (LaFayette and Travis 1990; Sticher *et al.* 1990; Helm *et al.* 1993). A subfamily of three endoplasmic reticulum intronless genes, Solyc01g102960, Solyc11g020330 and Solyc03g113930, is present in the tomato genome. Solyc01g102960 is highly differentially expressed only during fruit ripening. On the other hand, Solyc11g020330 and Solyc03g113930are highly differentially expressed during both fruit ripening and heat-shock stress in leaves and microspores, suggesting their role as general-purpose stress responsive sHSP genes at the endoplasmic reticulum.

Ripening involves a massive structural change comparable to that experimented by plastids when exposed to environmental stress (Giovannoni 2001). In particular, the cytosol is an important place for the activity of sHSP gene products in tomato. Similarly to other higher plants, the subfamily of cytosolic sHSP genes in tomato is larger than those of cell organelles (Reddy *et al.* 2014). Eight sHSP genes, conforming cytosolic subfamilies CI and CII, are highly differentially expressed during fruit ripening (Table S5). In agreement with the ancient membrecy of cytosolic genes to the sHSP gene family (Waters and Vierling 1999),these genes account for ~77% of the total transcripts of sHSP genes in the MR ripening stage (Table S2). Remarkably, intronless Solyc06g076560, the youngest sHSP gene exclusive to the *S. lycopersicum* clade accounting for ~17% of the total transcripts of sHSP genes in the MR ripening stage, is also highly differentially expressed during heat shock stress in leaves and microspores. In summary, Solyc03g082420, Solyc05g014280, Solyc11g020330, and Solyc06g076560, targeted to the CP, the ER and cytosolic compartments, define a core set of sHSP genes contributing to cell homeostasis in both fruit ripening and heat shock stress, suggesting that other subsets of multi-purpose sHSP genes may coexist within the family.

The results presented here for the sHSP gene family suggest that systematic approaches built upon an evolutionary analysis of transcriptome data may be also effective todisentangle the organization and functionality of other complex gene families.

## ACKNOWLEDGMENTS

This work was funded by project PICT 2012-2513, “Multiplex systems for targeted microfluidic amplification and NGS sequencing,” National Agency for Science and Technology Promotion, Argentina. Institution: National Scientific and Technical Research Council (CONICET) and PICT 2014-3181, “Process optimization of genetic breeding in tomato using bioinformatics tools”. National Agency for Science and Technology Promotion, Argentina. Institution: National Scientific and Technical Resea chCouncil (CONICET).

## LITERATURE CITED

Aanes, H., C. Winata, L. F. Moen, O. Østrup, S. Mathavan, P. Collas, T. Rognes, and P. Alestrom, 2014 Normalizationof rna-sequencing data from samples with varying mRNA levels. PloS one9: e89158.

Alba, R., P. Payton, Z. Fei, R. McQuinn, P. Debbie, G. B. Martin, S. D. Tanksley, and J. J. Giovannoni, 2005 Transcriptome and selected metabolite analyses reveal multiple points of ethylene control during tomato fruit development. The Plant cell 17: 2954–2965.

Andolfo, G., F. Jupe, K. Witek, G. J. Etherington, M. R. Ercolano, and J. D. G. Jones, 2014 Defining the full tomato NB-LRR resistance gene repertoire using genomic and cDNA RenSeq. BMC Plant Biology 14: 120.

Aoki, K., Y. Ogata, K. Igarashi, K. Yano, H. Nagasaki, E. Kaminuma, and A. Toyoda, 2013 Functional genomics of tomato in a post-genome-sequencing phase. Breeding Science 63: 14–20.

Baniwal, S. K., K. Bharti, K. Y. Chan, M. Fauth, A. Ganguli, S. Kotak, S.K. Mishra, L. Nover, M. Port, K.-D. Scharf, J. Tripp, C. Weber, D. Zielinski, and P. von Koskull-Doring 2004 Heat stress responsein plants:a complex game with chaperones and more than twenty heat stress transcription factors. Journal of Biosciences29: 471–487.

Basha, E., H., O’Neill, and E. Vierling, 2015 Small heat shock proteins and α-crystallins: dynamic proteins with flexible functions. Trends in Biochemical Sciences37: 106–117.

Bondino, H., E., Valle, and A. ten Have, 2012 Evolution and functional diversification of the small heat shock protein alpha-crystallin family in higher plants. Planta 235: 1299–1313.

Cannon, S. B., A. Mitra, A. Baumgarten, N. D. Young, and G. May, 2004 The roles of segmental and tandem gene duplication in the evolution of large gene families in Arabidopsis thaliana. BMC Plant Biology 4: 1–21.

de Jong, W. W., G. J. Caspers, and J. A. Leunissen, 1998 Genealogy of the alpha-crystallin-small heat-shock protein superfamily. International journal of biological macromolecules 22: 151–162.

Eisen, M. B., P. T. Spellman, P. O. Brown, and D. Botstein, 1998 Cluster analysis and display of genome-wide expression patterns. Proc Natl Acad Sci USA 95.

Emanuelsson, O., H. Nielsen, S. Brunak, and G. von Heijne, 2000 Predicting subcellular localization of proteins based on their n-terminal amino acid sequence. Journal of Molecular Biology 300: 1005 – 1016.

Ercolano, M. R., A. Sacco, F. Ferriello, R. D’Alessandro, P. Tononi, A. Traini, A. Barone, E. Zago, M. L. Chiusano, G. Buson, M. Delledonne, and L. Frusciante, 2014 Patchwork sequencing of tomato San Marzano and Vesuviano varieties highlights genome-wide variations. BMC genomics 15: 138.

Exposito-Rodriguez, M., A. Borges, A. Borges-Perez, and J. Perez, 2008 Selection of internal control genes for quantitative real-time RT-PCR studies during tomato development process. BMC Plant Biology 8: 131.

Faurobert, M., C. Mihr, N. Bertin, T. Pawlowski, L. Negroni, N. Sommerer, and M. Causse, 2007 Majorproteome variations associated with cherry tomato pericarp development and ripening. Plant physiology 143: 1327–46.

Fei, Z., X. Tang, R. M. Alba, J. a. White, C. M. Ronning, G. B. Martin, S. D. Tanksley, and J. J. Giovannoni, 2004 Comprehensive EST analysis of tomato and comparative genomics of fruit ripening. The Plant journal: for cell and molecular biology 40: 47–59.

Fragkostefanakis, S., S. Simm, P. Paul, D. Bublak, K.-D. Scharf, and E. Schleiff, 2015 Chaperone network composition in *Solanum lycopersicum* explored by transcriptome profiling and microarray meta-analysis. Plant, cell & environment 38: 693–709.

Frank, G., E. Pressman, R. Ophir, L. Althan, R. Shaked, M. Freedman, S. Shen, and N. Firon, 2009 Transcriptional profiling of maturing tomato (Solanum lycopersicumx L.) microspores reveals the involvement of heat shock proteins, ROS scavengers, hormones, and sugars in the heat stress response. Journal of Experimental Botany 60: 3891–3908.

Giovannoni, J., 2001 Molecular Biology of Fruit Maturation and Ripening. Annual review of plant physiology and plant molecular biology 52: 725–749.

Gonzalez-Porta, M., A. Frankish, J. Rung, J. Harrow, and A. Brazma, 2013 Transcriptome analysis of human tissues and cell lines reveals one dominant transcript per gene. Genome Biology 14: R70.

Goyal, R. K., V. Kumar, V. Shukla, R. Mattoo, Y. Liu, S. Chung, J. J. Giovannoni, andA. K. Mattoo, 2012 Features of a unique intronless cluster of class I small heat shock protein genes in tandem with box C/D snoRNA genes on chromosome 6 in tomato (Solanum lycopersicum). Planta 235: 453–471.

He, X. and J. Zhang, 2005 Rapid subfunctionalization accompanied by prolonged and substantial neofunctionalization in duplicate gene evolution. Genetics 169: 1157–1164.

Hebenstreit, D., M. Fang, M. Gu, V. Charoensawan, A. van Oudenaarden, and S. A. Teichmann, 2011 RNA sequencingreveals two major classes of gene expression levels in metazoan cells. Molecular Systems Biology 7: 497.

Helm, K. W., P.R. LaFayette, R.T. Nagao, J. L. Key, and E. Vierling, 1993 Localization of small heat shock proteins to the higher plant endomembrane system. Molecular and Cellular Biology 13:238–247.

Hicks, S.C. and R.A. Irizarry, 2015 quantro: a data-driven approach to guide the choice of an appropriate normalization method. Genome Biology 16: 1–8.

Hillis, D. M. and J. J. Bull, 1993 An Empirical Test of Bootstrapping as a Method for Assessing Confidence in Phylogenetic Analysis. Systematic Biology 42: 182–192.

Howell, S. H., 2013 Endoplasmic reticulum stress responses in plants. Annual Review of Plant Biology 64: 477–499, PMID: 23330794.

Jarvis, P. and E. López-Juez, 2013 Biogenesis and homeostasis of chloroplasts and other plastids. Nat Rev Mol Cell Biol 14: 787–802.

Kelley, L. A., S. Mezulis, C. M. Yates, M. N. Wass, and M.J. E. Sternberg, 2015 The Phyre2 web portal for protein modeling, prediction and analysis. Nat. Protocols 10: 845–858.

Krsticevic, F., H. Santos, S. Januário, C. Schrago, and A. Carvalho, 2010 Functional Copies of the Mst77F Gene on the Y Chromosome of Drosophila melanogaster. Genetics 184.

LaFayette, P. and R. Travis, 1990 Soluble and membrane-associated heat shock proteins in soybean root. Protoplasma 156: 174–182.

Lambert, W., P.J. Koeck, E. Ahrman, P. Purhonen, K. Cheng, D. Elmlund, H. Hebert, and C. Emanuelsson, 2011 Subunit arrangement in the dodecameric chloroplast small heat shock protein hsp21. Protein Science 20: 291–301.

Larkin, M. A., G. Blackshields, N. Brown, R. Chenna, P. A. McGettigan, H. McWilliam, F. Valentin, I. M. Wallace, A. Wilm, R. Lopez, et al., 2007 Clustal W and Clustal X version 2.0. Bioinformatics 23: 2947–2948.

Lawrence, S., K. Cline, and G. Moore, 1997 Chromoplast development in ripening tomato fruit: identification of cdnas for chromoplast-targeted proteins and characterization of a cdna encoding a plastid-localized low-molecular-weight heat shock protein. Plant Molecular Biology 33: 483–492.

Lee, J. M., J.-G. Joung R. McQuinn, M.-Y. Chung, Z. Fei, D. Tieman, H. Klee, and J. Giovannoni, 2012 Combined transcriptome, genetic diversity and metabolite profiling in tomato fruit reveals that the ethylene response factor slerf6 plays animportant role in ripening and carotenoid accumulation. The Plant Journal 70: 191–204.

Librado, P. and J. Rozas, 2009 DnaSP v5: a software for comprehensive analysis of DNA polymorphism data. Bioinformatics 25: 1451–1452.

Lopes-Caitar, V. S., M. C. de Carvalho, L. M. Darben, M. K. Kuwahara, A. L. Nepomuceno, W. P. Dias, R. V. Abdelnoor, and F. C. Marcelino-Guimarães, 2013 Genome-wide analysis of the hsp20 gene family in soybean: comprehensivesequence, genomic organization and expression profile analysis under abiotic and biotic stresses. BMC Genomics 14: 1–17.

Low, D., K. Brandle, L. Nover, and C. Forreiter, 2000 Cytosolic heat-stress proteins Hsp17.7 class I and Hsp17.3classII of tomato act as molecular chaperones in vivo. Planta 211: 575–582.

Lynch, M. andA. Force, 2000 The probability of duplicate gene preservation by subfunctionalization. Genetics 154.

MacCarthy, T. and A. Bergman, 2007 The limits of subfunctionalization. BMC Evolutionary Biology 7: 1–14.

Matas, A. J., T. H. Yeats, G. J. Buda, Y. Zheng, S. Chatterjee, T. Tohge, L. Ponnala, A. Adato, A. Aharoni, R. Stark, A. R. Fernie, Z. Fei, J. J. Giovannoni, andJ. K. Rose, 2011 Tissue-and cell-type specific transcriptome profiling of expanding tomato fruit provides insights into metabolic and regulatory specialization and cuticle formation. The Plant Cell 23: 3893–3910.

Montgomerie, S., J. A. Cruz, S. Shrivastava, D. Arndt, M. Ber-janskii, and D. S. Wishart, 2008 Proteus2: a web server for comprehensive protein structure prediction and structure-based annotation. Nucleic Acids Research 36: W202–W209.

Neta-Sharir, I., T. Isaacson, S. Lurie, and D. Weiss, 2005 Dual Role for Tomato Heat Shock Protein 21: Protecting Photosystem II from Oxidative Stress and Promoting Color Changes during Fruit Maturation. The Plant Cell 17: 1829–1838.

Ohno, S., 1970 Evolution by Gene Duplication. Springer Berlin Heidelberg, Berlin, Heidelberg.

Ouyang, Y., J. Chen, W. Xie, L. Wang, and Q. Zhang, 2009 Comprehensivesequence and expression profile analysis of hsp20 gene family in rice. Plant Mol Biol 70.

Poulain, P., J.-C. Gelly, andD. Flatters, 2010 Detection and architecture of small heat shock protein monomers. PloS one 5: e9990.

Prasinos, C., K. Krampis, D. Samakovli, andP. Hatzopoulos, 2005 Tight regulation of expression of two Arabidopsis cytosolic Hsp90 genes during embryo development. Journal of experimental botany 56: 633–44.

Ramakrishna, W., Z. Deng, C. K. Ding, A. K. Handa, and R. H. Ozminkowski, 2003 A novel small heat shock protein gene, vis1, contributes to pectin depolymerization and juice viscosity in tomato fruit. Plant Physiol. 131: 725735.

Reddy, P. S., P. B. Kavi Kishor, C. Seiler, M. Kuhlmann, L. Eschen-Lippold, J. Lee, M. K. Reddy, and N. Sreenivasulu, 2014 Unraveling regulation of the small heat shock proteins by the heat shock factor hvhsfb2c in barley: Its implications in drought stress response and seed development. PLoS ONE 9: e89125.

Riechmann, J. L., J. Heard, G. Martin, L. Reuber, C.-Z. Jiang, J. Keddie, L. Adam, O. Pineda, O. J. Ratcliffe, R. R. Samaha, R. Creelman, M. Pilgrim, P. Broun, J. Z. Zhang, D. Ghandehari, B. K. Sherman, and G. L. Yu, 2000 Arabidopsis transcription factors: Genome-wide comparative analysis among eukaryotes. Science 290: 2105–2110.

Robinson, M. D., D. J. McCarthy, and G. K. Smyth, 2010 edgeR: a Bioconductor package for differential expression analysis of digital gene expression data. Bioinformatics 26: 139–140.

Sanmiya, K., K. Suzuki, Y. Egawa, and M. Shono, 2004 Mitochondrial small heat-shock protein enhances thermotolerance in tobacco plants. FEBS Letters 557: 265–268.

Scharf, K. D., M. Siddique, and E. Vierling, 2001 The expanding family of Arabidopsis thaliana small heat stress proteins and a new family of proteins containing alpha-crystallin domains (acd proteins). Cell Stress and Chaperones 6.

Siddique, M., S. Gernhard, P. von Koskull-DÖring, E. Vierling, and K.-D. Scharf, 2008 The plant shsp superfamily: five new members in Arabidopsis thaliana with unexpected properties. Cell Stress and Chaperones 13: 183–197.

Srivastava, A., A. Gupta, D. t, M. Ak, and H. Ak, 2010 Maturity and ripening-stage specific modulation of tomato (Solanum lycopersicum) fruit transcriptome. GM Crops 1: 237–249., PMID: 21844679.

Sticher, L., A. K. Biswas, D. S. Bush, and R. L. Jones, 1990 Heat Shock Inhibits *α*-Amylase Synthesis in Barley Aleurone without Inhibiting the Activity of Endoplasmic Reticulum Marker Enzymes. Plant Physiology 92: 506–513.

Tamura, K., G. Stecher, D. Peterson, A. Filipski, and S. Kumar, 2013 Mega6: Molecular evolutionary genetics analysis version 6.0. Molecular Biology and Evolution 30: 27252729.

Tang, H., J. E. Bowers, X. Wang, R. Ming, M. Alam, and A. H. Paterson, 2008 Synteny and collinearity in plant genomes. Science 320: 486–488.

The Tomato Genome Consortium, 2012 The tomato genome sequence provides insights into fleshy fruit evolution. Nature 485: 635–41.

Trapnell, C., L. Pachter, and S. L. Salzberg, 2009 TopHat: discovering splice junctions with RNA-Seq. Bioinformatics (Oxford, England) 25: 1105–11.

Walter, P. andD. Ron, 2011 The unfolded protein response:From stress pathway to homeostatic regulation. Science 334: 1081–1086.

Waters, E. R., 1995 The Molecular Evolution of the Small Heat-Shock Proteins in Plants. Genetics 141: 785–95.

Waters, E. R., B. D. Aevermann, andZ. Sanders-Reed, 2008 Comparative analysis of the small heat shock proteins in three angiosperm genomes identifies new subfamilies and reveals diverse evolutionary patterns. Cell Stress and Chaperones13.

Waters, E. R., G. J. Lee, and E. Vierling, 1996 Review article: evolution, structure and function of the small heat shock proteins in plants. J Exp Bot 47.

WatersE. R. andE. Vierling, 1999 The diversification of plant cytosolic small heat shock proteins preceded the divergence of mosses. Molecular Biology and Evolution 16: 127–139.

Xia, K., T. Liu, J. Ouyang, R. Wang, T. Fan, and M. Zhang, 2011 Genome-wide identification, classification, and expression analysis of autophagy-associated gene homologues in rice (oryza sativa l.). DNA research p. dsr024.

Yang, Z., Y. Zhou, X. Wang, S. Gu, J. Yu, G. Liang, C. Yan, and C. Xu, 2008. Genomewide comparative phylogenetic and molecular evolutionary analysis of tubby-like protein family in arabidopsis, rice, and poplar. Genomics 92: 246–253.

Zeng, Y., Z. Pan, L. Wang, Y. Ding, Q. Xu, S. Xiao, and X. Deng, 2014 Phosphoproteomic analysis of chromoplasts from sweet orange during fruit ripening. Physiologia Plantarum 150: 252–270.

Zhao, C., M. Shono, A. Sun, S. Yi, M. Li, and J. Liu, 2007 Constitutive expression of an endoplasmic reticulum small heat shock protein alleviates endoplasmic reticulum stress in transgenic tomato. Journal of Plant Physiology 164: 835–841.

Zheng, D. and M. B. Gerstein, 2007 The ambiguous boundary between genes and pseudogenes: the dead rise up, or do they? Trends in Genetics 23: 219 – 224.

